# Integrating Amaranth and Agrivoltaics in the Creation and Assessment of Climate Resilient Food Systems

**DOI:** 10.64898/2026.06.19.732940

**Authors:** Jarvinia Rowe-Ibekwe, Uzair Jamil, Joshua M. Pearce, Raymond Thomas, Linda Alrayes

## Abstract

Food insecurity affects 8.69M Canadians and is projected to increase due to climate change. Amaranth, a resilient crop with high nutrient content, is an ideal candidate for building climate-resilient food systems. This study investigates whether amaranth grown under different solar photovoltaic (PV) modules can be used to develop and assess climate-resilient food systems under simulated present and future climates using sealed biomes. We found increased vertical growth, leaf temperature, and production under simulated 2050 climate, but reduced photosynthesis, transpiration, CO2 uptake, and stomatal conductance by 9.4%, 51.2%, 48%, and 50%, respectively, compared to 2024 plants. Crops grown under 69% transparent crystalline silicon-based PV modules restored these physiological performances by 4.7%, 72.9%, 5.4%, and 100%, respectively, compared to 2025 plants without agrivoltaics. This research suggests integrating amaranth with agrivoltaics as a viable strategy to produce climate-resilient food systems that enhance food security, sustainable energy, and economic stability under changing climates.

## Main

Food insecurity is a pressing issue in Canada, particularly in rural, coastal and minority communities. According to the 2022 Canadian Income Survey, approximately 8.69 million Canadians lived in food-insecure households, a 4.5% increase from the previous year^1^. Additionally, climate change will complicate food production and accessibility by threatening crop performance and arable land from more extreme and unpredictable weather patterns^2^. To meet the nutritional needs of a growing population^3^ while curbing food insecurity trends, innovative and climate resilient agricultural practices that also support ecological and economic sustainability are urgently needed^4^.

Agrivoltaics is the dual use of land for the coproduction of agriculture and electricity, offering a promising solution to sustainable and climate-resilient food systems. It enhances crop yields ^5–9^, reduces greenhouse gas emissions ^10^ and improves economic and environmental sustainability ^11^. The technology can improve water consumption ^12^, protect crops from high winds ^13^, and reduce soil erosion^14^. With the integration of agrivoltaics, complete transformation of deserted areas has been reported ^15^. Agrivoltaics also creates microclimates that enhance plant growth, and conserve ecosystem biodiversity ^16^. Studies in Canada have shown immense agrivoltaics potential and almost a third of Canada’s current electricity requirements could be catered by converting 1% of agricultural land to agrivoltaics^10^. Moreover, initial trials with lettuce ^17,18^ and strawberries ^19,20^ have shown encouraging results.

*Amaranthus Viridis* is a nutrient-dense, fast-growing leafy vegetable ^21^. It contains more protein than common cereals like corn, wheat, and rice and surpasses other leafy vegetables such as kale and spinach in nutritional value ^22^. Amaranth can also thrive in saline, dry or nutrient-poor soils and withstand extreme temperatures ^23,24^, making it ideal for innovative climate-resilient food production systems like agrivoltaics. A recent study by Jamil et al. (2025) focused on the optimization of amaranth under agrivoltaics ^25^, however, it did not address underlying biological mechanisms behind plant performance in this food system. To fully reveal insights into amaranth resilience and productivity under agrivoltaics, its agronomic and physiological responses under different climatic conditions must be assessed.

Given the complexity of plant biological interactions, recording growth and physiological responses can generate large datasets, requiring advanced computational methods for analysis. Bioinformatics is essential for identifying relationships between plant growth and environmental or management factors ^26^. It offers a systems biology perspective to better understand interactions between growth performance indicators and physiological responses enabling data-driven decision support for innovating and assessing climate resilient agrivoltaics based production systems. Key growth performance indicators including yield, plant height, leaf number and leaf surface area are critical to biomass accumulation and photosynthesis ^27,28^. Physiological traits such as photosynthetic rate, CO_2_ uptake, stomatal conductance, photosynthesis, transpiration, chlorophyll fluorescence, leaf temperature and PAR play a significant role in explaining plant adaptation to stress ^29,30^. Hence, understanding these variables are essential for assessing and predicting growth and physiological response to changing climates in different crop management systems.

Despite amaranth being an emerging superfood crop with high demands in many locations globally, it is underutilized in temperate climate-based food production systems as a potential solution to food insecurity. Furthermore, its utility in building climate resilient, sustainable, food systems under emerging technologies such as agrivoltaics is unexplored. To address this gap, we assessed amaranth’s growth and physiological responses to varying climates when cultivated under different semi-transparent PV in simulated current (2024) and future (2050) climate conditions using sealed biomes.

This research may inform the optimal design of agrivoltaic systems for amaranth cultivation, highlighting their potential as a sustainable strategy to enhance food security and agricultural resilience in response to climate change for horticultural crop production ^31^. The findings could optimize the balance between low and high PV shading for amaranth, enhancing resilience and productivity under changing environmental conditions ^32^. Output for this work may also inform system design and policy, particularly in boreal regions facing food insecurity, and aligns with the UN Sustainable Development Goals, specifically Zero Hunger, Affordable and Clean Energy and Climate Action ^33^.

## Results

### Analysis 1: How Will Climate Change Impact Amaranth Physiological and Growth Performance?

#### Comparing Amaranth Growth and Physiology when Cultivated Under Current and Future Climate Conditions

The physiology and growth of amaranth without agrivoltaic intervention were analysed. In comparison to plants grown in current climate, those grown in 2050 climate conditions exhibited greater height of 20.4 cm across all five weeks of growth and harvest periods (Figs. 2C & 2D). Plant yield in both climates fluctuated over the five weeks of harvest (Fig. 1).

**Figure 1.**
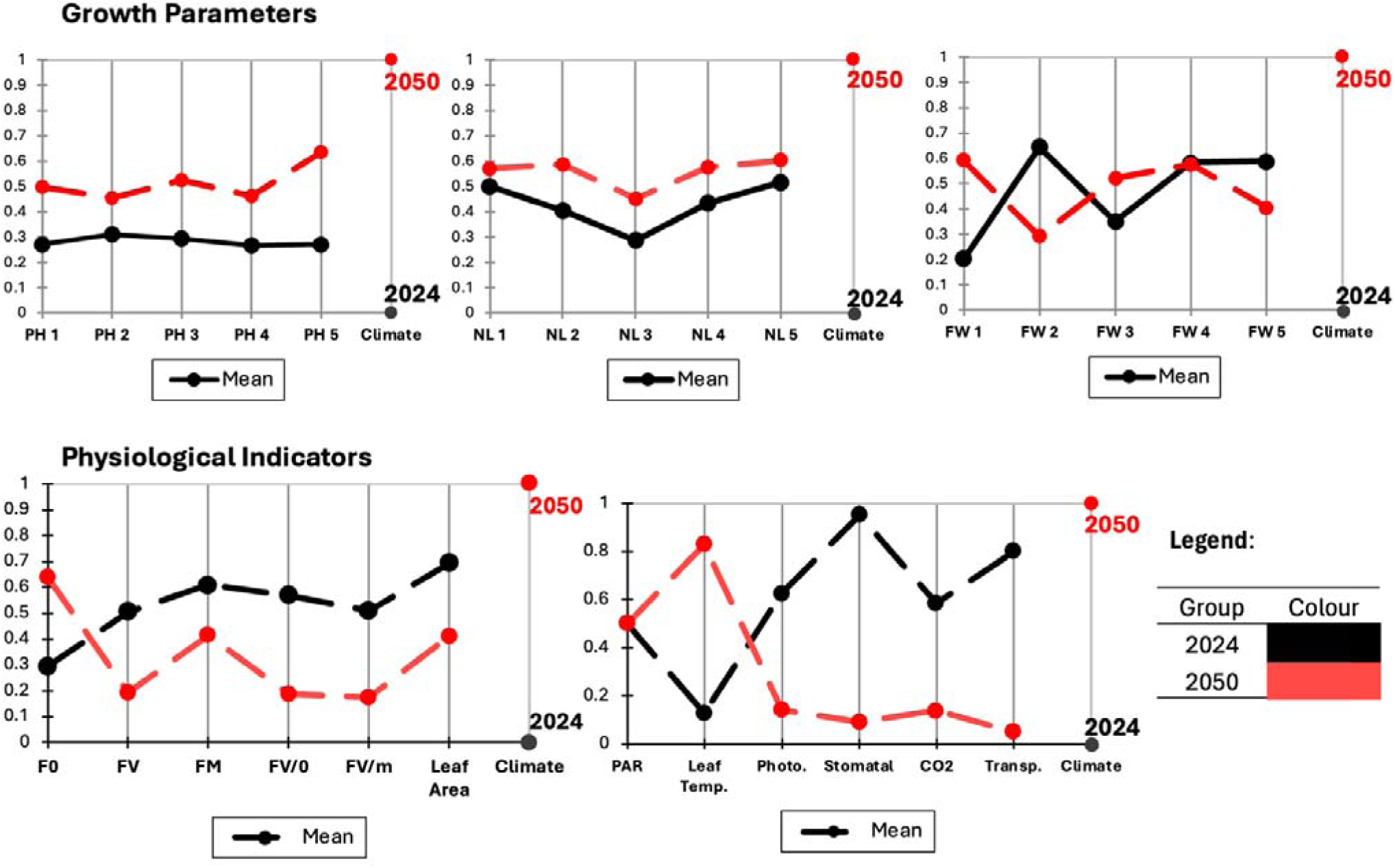
Parallel plots illustrating relationships between amaranth growth parameters (yield, plant height, leaf number) and physiological traits (photosynthetic rate, stomatal conductance, CO_2_ uptake, transpiration and PAR) across simulated 2024 and 2050 climate conditions. The spread of the data is represented by the rescaled mean for n = 8 plant replicates per climate treatment. Measured variables (y-axis) with averaged data points for 2024 (black line) and 2050 (red line) treatments.

**Figure 2.**
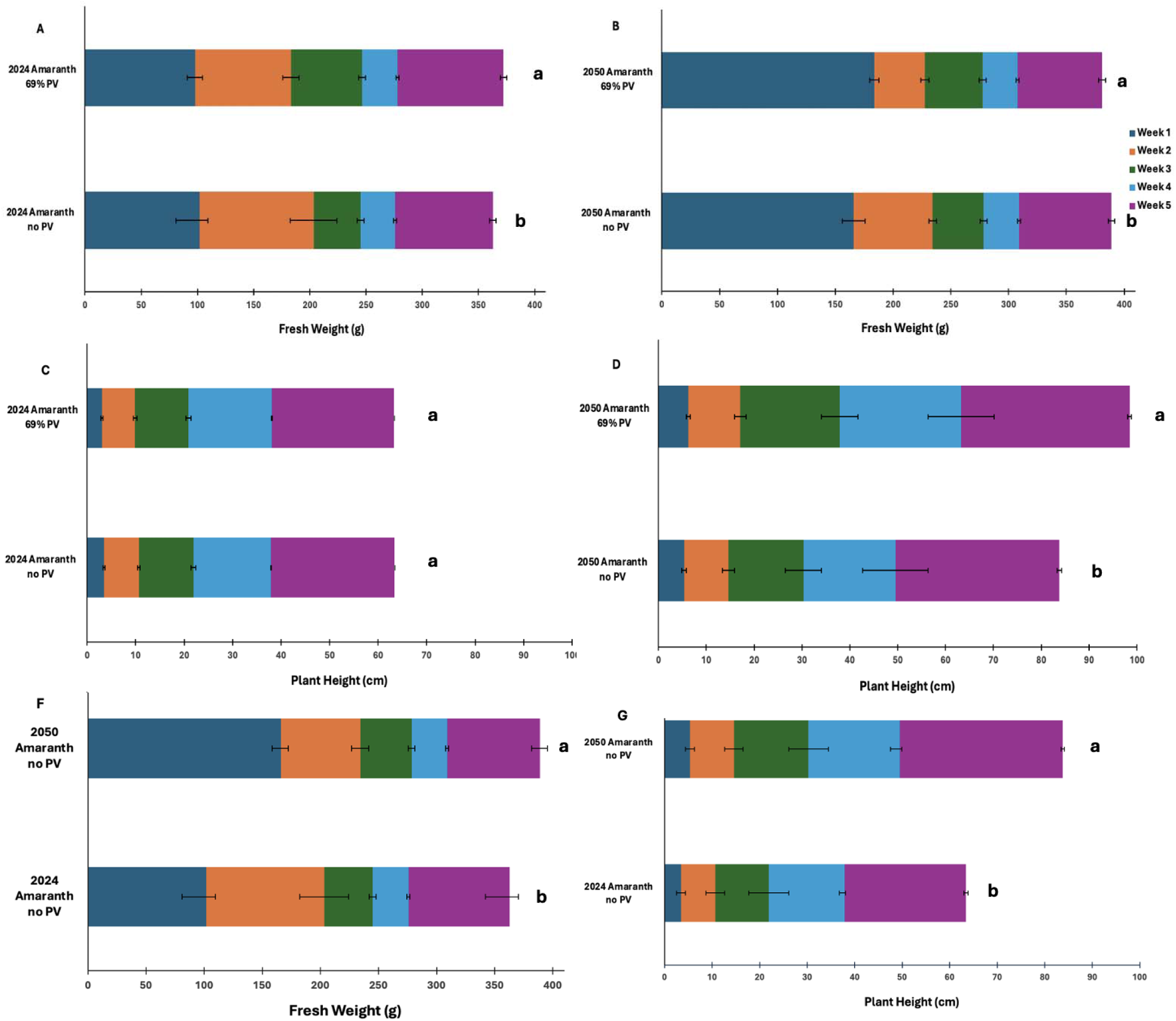
Bar chart illustrating relationships between amaranth harvest (A, B, F) and vertical growth (C, D, G) across simulated 2024 and 2050 climate conditions with and without agrivoltaic intervention. Represented are the pooled average yield and height ± standard error for n = 8 plant replicates per climate treatment. The weight of each harvest and the height of the plant (x-axis) are recorded in grams and centimetres respectively with weekly harvests distinguished by different colours.

When comparing plants without agrivoltaic treatment, the leaf area, stomatal conductance, intracellular CO_2_, photosynthetic rate and leaf transpiration were higher in amaranth grown in 2024 conditions. In addition, all chlorophyll and photosynthesis-related physiological parameters, except for F0 values, were higher in amaranth grown under 2024 conditions. PAR values were similar between plants grown in both climates without photovoltaic treatment.

#### Applying different multivariate analysis to determine how climate change impacted amaranth growth and physiology

Among the multivariate methods evaluated, partial least squares discriminant analysis (PLS-DA) was the most effective technique for clustering and ordinating growth and physiological variables based on climate conditions. The variable importance in projection (VIP) scores, identified the most important growth and physiological variables that explained the differences in crop performance between the two climate conditions (Fig. 3C). Variables like transpiration, stomatal conductance, leaf temperature, photosynthesis, intracellular CO_2_, fresh weight in weeks one and two, plant height in week five of growth, minimal fluorescence, variable fluorescence and initial fluorescence response with VIP scores above 1.0 contributed the most to the differentiation between the two climate conditions. Variables with VIP scores below 1.0 indicated lower differentiation and importance with respect to the two climate conditions.

**Figure 3.**
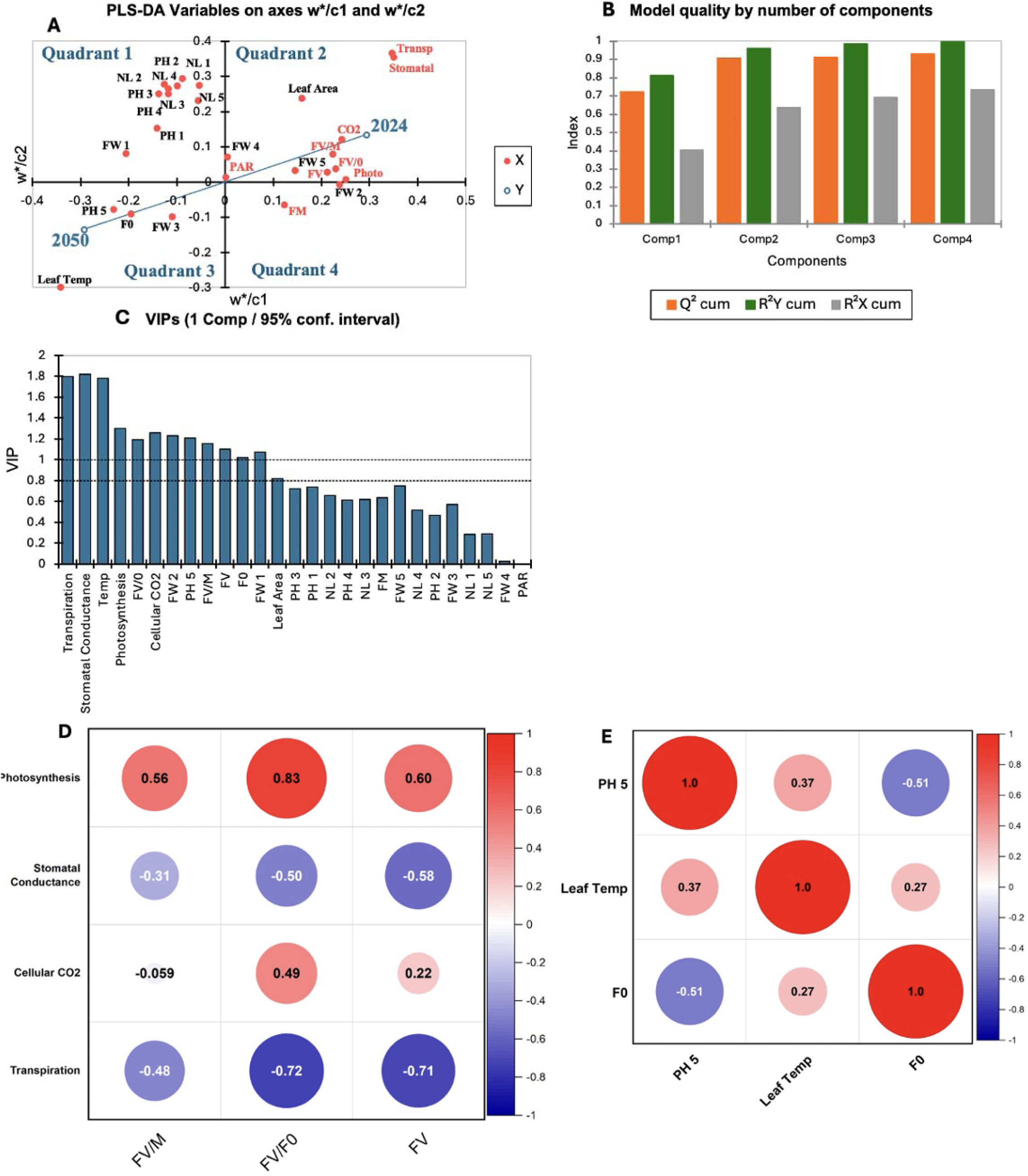
Partial least square discriminant analysis (PLS-DA) and correlation analyses of physiological and growth responses of amaranth under simulated current (2024) and future (2050) climate conditions in sealed biomes. PLS-DA analysis was used to evaluate variable clustering based on climate conditions of n = 8 plants replicates per treatment, as it provided the most reliable projections (A). The model’s quality is supported by Q^2^ scores for four components: 0.722, 0.908, 0.911 to 0.931indicating high model quality (B). The VIP scores from the analysis highlight variables that contribute to the model (C). Variables with a VIP score below 1 are not considered significant in clustering between groups, while those with scores above 1 are highly influential. The correlation plot displays the significant Pearson correlation coefficients between photosynthesis, stomatal conductance, intracellular CO_2_, transpiration and FV, FV/F0 and FV/FM of amaranth cultivated under 2024 climatic conditions (D). Under 2050 climate conditions, the correlation plot displays the significant correlation coefficients between PH 5, Leaf Temp, and F0 (E).

PLS-DA results showed a clear ordination of growth and physiological variables that were associated with each climate condition. Quadrant two of the PLS-DA distinctly grouped leaf area, fresh weight (week five), photosynthesis, stomatal conductance, intracellular CO_2_, transpiration, variable fluorescence, initial fluorescence response and photosynthetic efficiency, which were all strongly associated with 2024 climate conditions (Fig. 3A).

In contrast, quadrant three of the PLS-DA clustered plant height (week five), minimum fluorescence, leaf temperature and fresh weight (week three), which were aligned with 2050 climate conditions (Fig. 3A). Other variables did not show clear ordination with current or future climates (Fig. 3A).

The PLS-DA is characterized by four components, representing the axes of data projection (Fig. 3B). The model exhibited strong predictive performance with Q^2^cumulative values increasing steadily across the four components: 0.722, 0.908, 0.911 to 0.931. This indicates strong predictive power as values above 0.8 demonstrate strong model accuracy. R²Y cumulative reflects the explained variance in the dependent variables which increased from 0.813 to 0.997 in all four components, demonstrating the model’s high ability in capturing the variance in explanatory (dependent) variables. Meanwhile, R^2^Xcumulative values increased from 0.406 (in component one) to 0.7 (in component four), indicating robust capacity to capture variance in the independent variables, confirming that PLS-DA is well-suited for analysing growth and physiological data trends in response to climate variation.

Other analyses like Factor Analysis, PCA, k-means clustering, MCA, AHC and RDA were used, however they were not the most suitable showing the relationship between the growth and physiological response (Supplementary Figs. 1 & 2).

#### Correlating Physiological Response and Growth Performance of Amaranth When Cultivated in Current (2024) Climate

ANOVA determined significant physiological and growth variables related to amaranth under 2024 climate conditions in quadrant two of the PLSDA model biplot.

Pearson’s correlation analysis highlighted the relationships between any significant physiological and growth variables that were related to 2024 climate (Fig. 3D). Photosynthesis showed a strong positive correlation with the initial fluorescence response (r = 0.83). Stomatal conductance exhibited a strong negative correlation with variable fluorescence (r = −0.58). Transpiration of amaranth cultivated in 2024 climate showed a strong negative correlation with the initial fluorescence response (r = −0.72) and variable fluorescence (r = −0.71). Transpiration and the photosynthetic efficiency exhibited a moderate negative correlation (r = −0.48). Photosynthesis also had a moderate positive correlation with variable fluorescence (r = 0.60) and photosynthetic efficiency (r = 0.56). Stomatal conductance had a moderate positive correlation with the initial fluorescence response (r = 0.50). The correlation between stomatal conductance and photosynthetic efficiency was moderately negative (r = −0.31). Intercellular CO_2_ had a moderate positive correlation with the initial fluorescence response (r = 0.49).

#### Relationship between Physiological Response and Growth Performance of Amaranth When Cultivated in Future (2050) Climate

ANOVA analysis determined significant physiological and growth variables ordinated with amaranth cultivated under 2050 climate conditions in quadrant three of the PLSDA model biplot. The growth and physiological variables of amaranth plants grown under simulated 2050 climate, were ordinated in quadrant three of the PLS-DA biplot (Fig. 3E) based on significant association. Pearson’s correlation analysis highlighted that height of amaranth plants in the fifth week of growth showed a strong negative correlation with the minimum fluorescence (r = −0.51). A moderate positive correlation was observed between plant height in the fifth week and leaf temperature (r = 0.37).

### Analysis 2: Can Agrivoltaics Mitigate the Impact of Climate Change on Amaranth Physiology and Growth Performance?

#### Assessing the Effects of Agrivotaics Production on Amaranth Growth Performance and Physiological Response under Future (2050) Climates

Amaranth cultivated under with 69% transparent crystalline silicon (c-Si) PV were taller throughout the entire cultivation cycle compared to control plants in 2050 (Fig. 4). They also developed more leaves than the control plants in weeks 2, 3, 4 and 5; however, they exhibited lower leaf surface area compared to the control plants during the growth period. Yield varied between the control and 69% PV plants. In terms of physiological response to 2050 climate, plants cultivated under 69% PV outperformed control plants in most measured parameters (Table S1).

**Figure 4.**
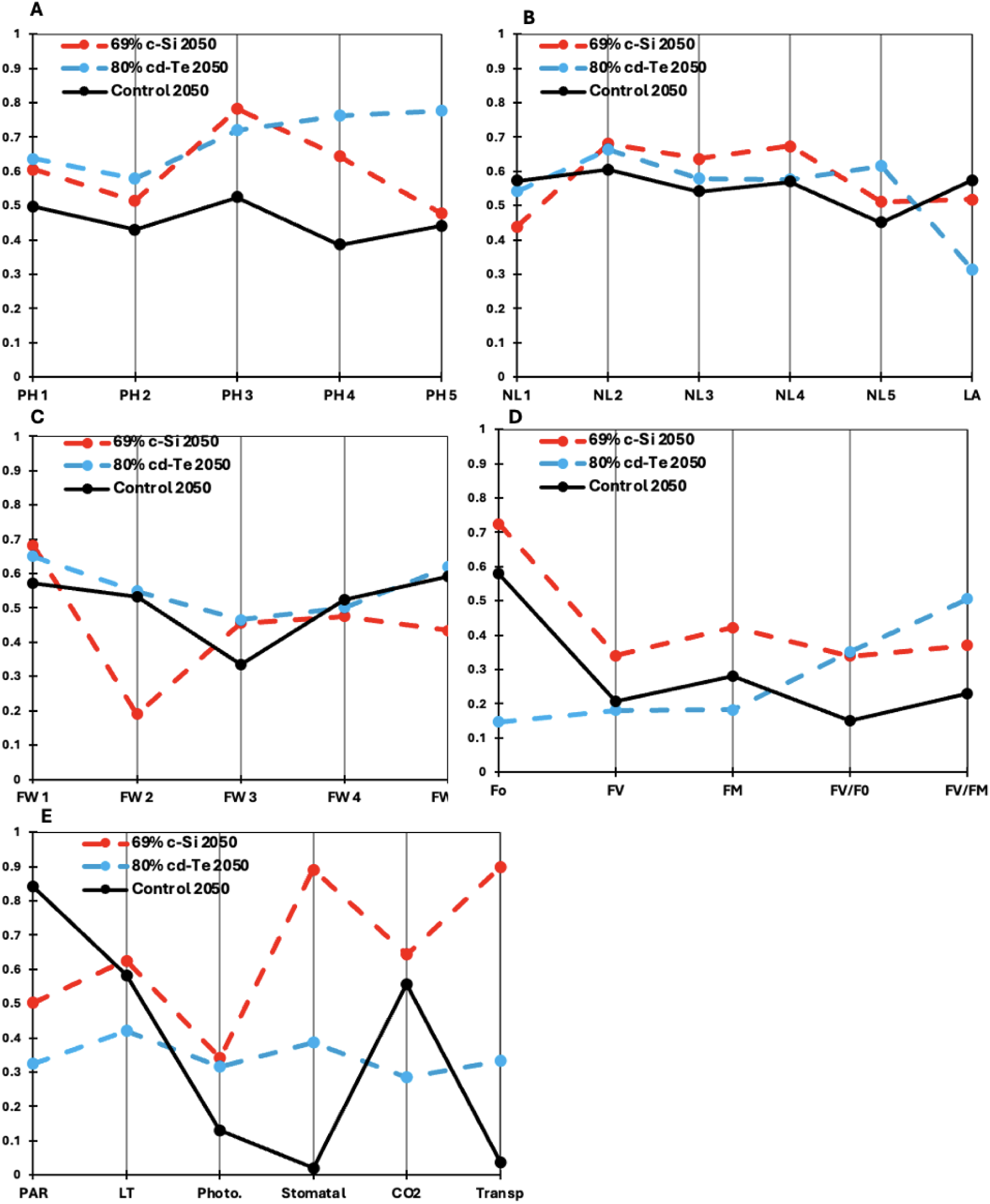
Parallel plots of the relationships between growth (A-C) and physiological (D,E) parameters of amaranth grown under control conditions, 69% crystalline silicon (c-Si) and 80% cadmium telluride (Cd-Te) photovoltaic cells in 2050 environmental conditions. The spread of the data is represented by the rescaled mean for n = 8 plant replicates per climate treatment. The y-axis axis denotes the measured variables, with data points for 69% c-Si, 80% Cd-Te and control treatment shown as red, blue and black lines, respectively.

The physiology and growth responses of amaranth were also compared with PV of different transparencies and materials (Fig. 4). Amaranth cultivated under 80% transparent cadmium telluride (Cd-Te) panels were taller than control plants. The plants grown under 80% transparent Cd-Te modules had more vegetative growth when compared to control plants in weeks 2, 3 and 5; however, the leaves had less surface area. Similar to plants under 69% transparent PV, yield also varied between the control and 80% transparent Cd-Te plants.

Of all the treatments, plants under 80% transparent Cd-Te modules had the highest photosynthetic efficiency. Physiologically, leaf temperature and intracellular CO□ levels were the lowest in 80% transparent Cd-Te plants and had lower photosynthetic rate, stomatal conductance and transpiration rate than 69% transparent c-Si amaranth.

#### Assessing Amaranth Performance under Agrivoltaics Production using Photovoltaics Panels with Varying Transparencies Across Current (2024) and Future (2050) Climate Scenarios

PLS-DA was the most effective technique for ordinating amaranth growth and physiological variables with specific PV transparencies under simulated 2024 and 2050 climate conditions (Figs. 5A-C).

**Figure 5.**
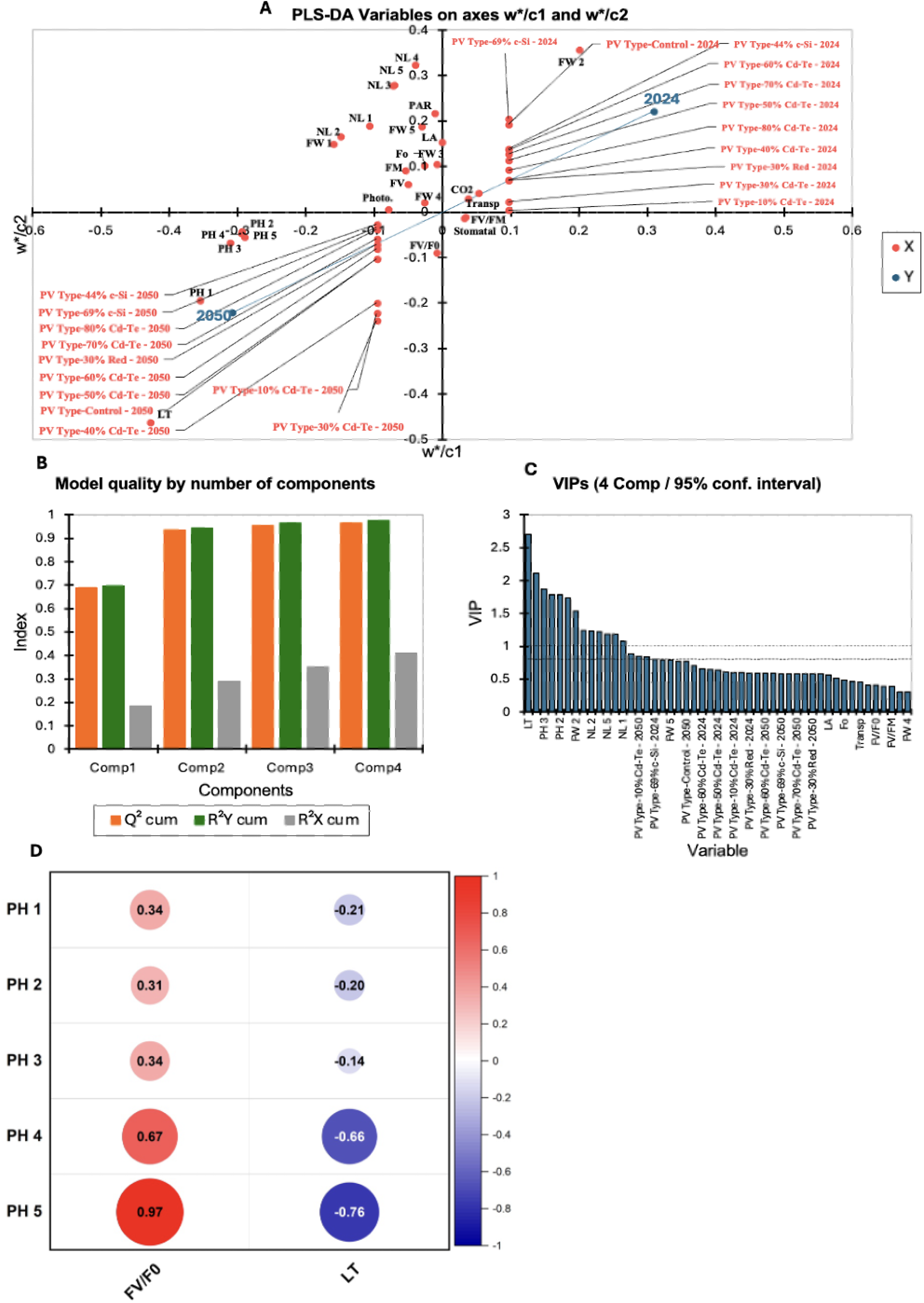
Multivariate and Pearson correlation analyses of physiological and growth responses of amaranth under a 2050 climate change scenario under 69% c-Si PV treatments. PLS-DA was chosen for further evaluation of variable clustering based on climate conditions, as it produced the most accurate variable projections (A). The Q2 scores for four components: 0.688, 0.935, 0.956 and 0.964 indicate a high-quality model (B). VIP scores from the analysis highlight which variables contribute to the model (C). Variables with a score below 1.0 are not considered influential in clustering between groups, while those above 1.0 play a significant role in differentiating groups. The heatmap shows the Pearson correlation coefficients between PH1-5 and Leaf Temp and FV/F0 (D).

The second quadrant of the PLS-DA indicated that intercellular CO_2_, transpiration and fresh weight in week two were clustered with 2024 climate conditions. These responses were predominantly associated with the 2024 climate scenario, particularly in plants grown without PV, as well as under 10%-80% transparent Cd-Te PV, 30% transparent Red PV and 44% and 69% transparent c-Si PV (Fig. 5A).

In the second quadrant, plant height, leaf temperature and initial fluorescence response were grouped with control plants (cultivated in 2050 with no PV), and those under 10%-80% transparent Cd-Te PV, 30% transparent Red PV and 44% and 69% transparent c-Si PV modules in 2050 climate conditions (Fig. 5A). Other variables did not cluster strongly with either climate condition in response to PV modules.

The PLS-DA model demonstrated high quality, capturing large amounts of variance in the dataset. Q^2^cumulative values for all components were all above 0.5, with scores of 0.688, 0.935, 0.956 and 0.964, respectively (Fig. 5B). The R^2^Xcumulative values increased from 0.185 in component one to 0.411 in component two, indicating a moderate explanation of variance within explanatory variables. Additionally, R^2^Ycumulative values increased from 0.698 in component one to 0.977 in component four, further supporting the model’s strong predictive power, although some variance in explanatory variables remains unaccounted for. Using VIP scores, the model identified the most influential variables in data ordination (Fig. 5C). The key contributors (VIP > 1.0) included leaf temperature, plant height in weeks two and three, fresh weight in week one and the number of leaves in weeks one, two and five.

#### Correlation of Physiological and Growth Variables in Amaranth under the Best-Performing PV

An ANOVA determined significant physiological and growth variables related to plants grown under 69% transparent c-Si modules in 2050 climate conditions in quadrant three of the PLSDA model biplot (Fig. 5A).

The relationship between statistically significant variables associated with the best-performing PV type in 2050 was analysed using Pearson correlation analysis (Fig. 5D). In amaranth cultivated under the 69% transparent c-Si PV, plant height during weeks 1-3 of growth showed a moderate positive correlation with variable fluorescence to initial fluorescence (r = 0.34, r = 0.31, and r = 0.34, respectively). In weeks four and five, this correlation became stronger (r = 0.67 and r = 0.97, respectively). In weeks 4 and 5 of plant height there was a negative correlation with leaf temperature strengthened (r = −0.66 and r = −0.76, respectively).

#### Economic Outlook

Agricultural revenues observed yield enhancements of 5.4% for 80% Cd-Te and 2.1% for 69% c-Si modules under projected 2050 and 2024 climate conditions (Fig. 6). Electrical revenues are calculated using initial electricity generation values for each module type, reduced annually by a 0.5% degradation rate. The capital cost of PV installation is assumed to be 2.16 CAD $/W, divided evenly over 50 years and subtracted from annual electrical revenue. Economic factors include a food price inflation rate of 3.9% per year and an electricity price inflation rate of 4.91%, consistent with historic Canadian trends. Amaranth price begins at 13.18 CAD $/kg, while the initial electricity sale price is 0.192 CAD $/kWh Across all years, total revenues from both agrivoltaic configurations substantially exceed traditional farming.

**Figure 6.**
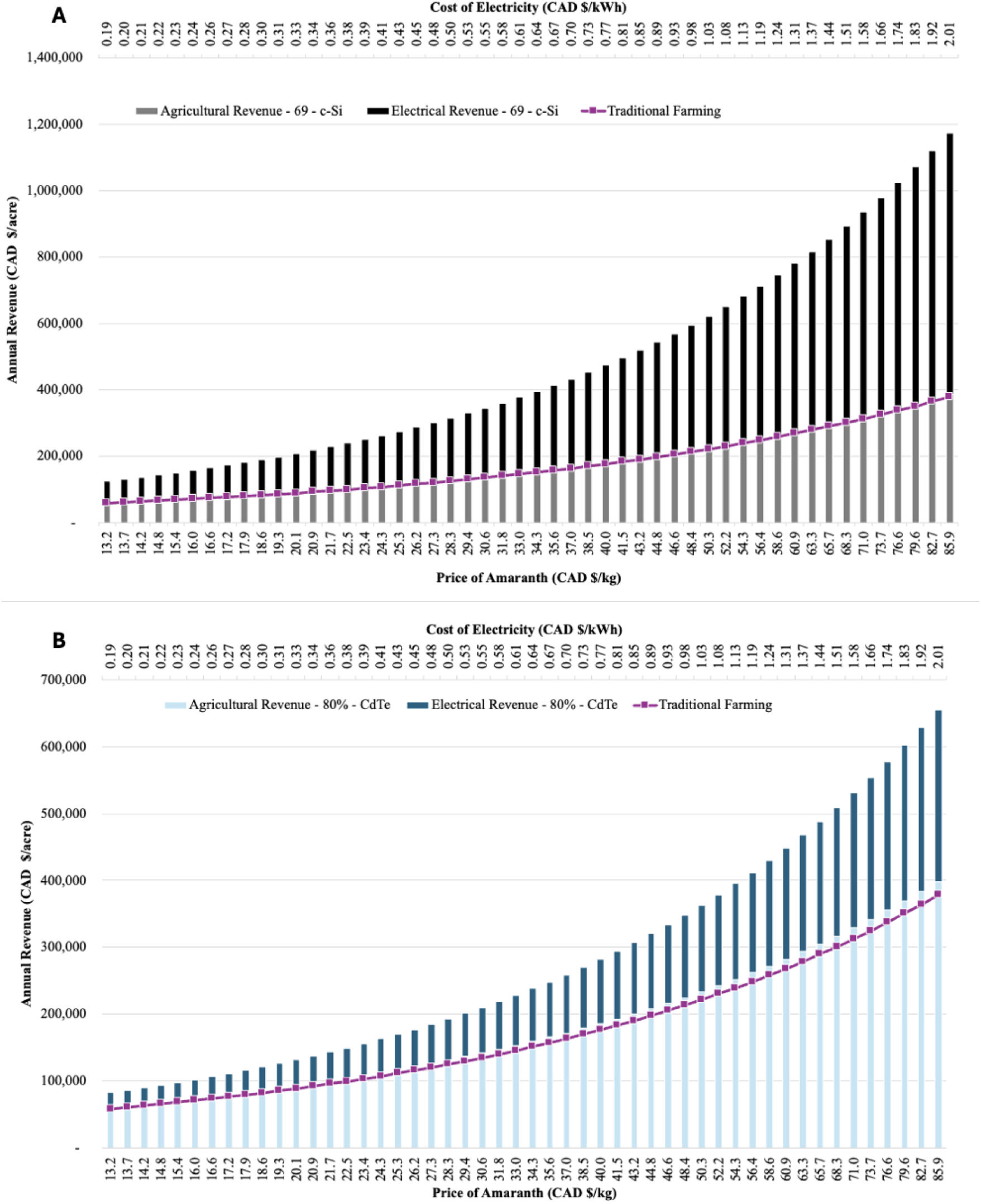
A 50-year comparative revenue analysis for 1 acre of amaranth farmland under three scenarios: agrivoltaics using 80% CdTe modules (A), agrivoltaics using 69% c-Si modules (B), and traditional farming without photovoltaics.

## Discussion

### Effect of Climate Change on Amaranth Growth Performance and Physiological Response

The impacts of climate change on amaranth were characterized by elevated leaf temperatures, along with reduced photosynthetic rates, transpiration, intercellular CO□, and stomatal conductance; clear signs of physiological stress (Fig. 4). In C4 plants like amaranth, increased temperature induces partial stomatal closure by increasing vapor pressure deficit, reducing water loss ^34^. This limits CO□ diffusion into intercellular airspaces, decreasing intercellular CO□ concentration. Prolonged thermal stress, such as the one simulated under 2050 climate scenarios, impairs photosynthetic efficiency due to reductions in Rubisco activity and enzyme stability. The combination of stomatal closure restricting CO□ influx and biochemical limitations constraining CO□ fixation reduces intercellular CO□ concentration and photosynthesis. This concurrent reduction in CO□ availability and assimilation reflects a breakdown between gas exchange and metabolic capacity, indicative of impaired photosynthetic regulation under elevated temperature stress. Although 2050 amaranth without PV exhibited increased plant height and leaf number compared to 2024 plants under similar treatment, this was not reflected in enhanced physiological performance or yield (Fig. 2). This discrepancy is due to a reduction in total leaf area, which limited light interception, canopy development, and biomass accumulation. The decline in photosynthetic function was not attributed to differences in PAR, which remained consistent across climate scenarios ^35^. Instead, elevated temperatures likely impaired Photosystem II efficiency, as indicated by decreased Fv/Fm and increased F□; markers of photoinhibition and thermal stress ^36^. This is consistent with findings showing that elevated leaf temperatures reduced intercellular CO□ concentration and impaired photosynthetic efficiency by disrupting gas exchange dynamics and enzymatic activity ^37^.

Although elevated atmospheric CO□ can enhance vegetative biomass without improving photosynthesis ^38^, this did not lead to increased yield under non-drought conditions. The absence of water limitation, combined with thermal and physiological stress, constrained overall productivity. Thus, while certain structural traits expanded in a future climate change scenario, impaired physiological function limited the yield potential of amaranth.

### Using PVs with varying transparencies to mitigate the adverse effects of climate change and innovate climate resilient food system

While Amaranth shows strong potential for innovating climate resilient food systems due to its ability to adapt to varying and harsh climate conditions ^23,24^, there is limited knowledge on its production under agrivoltaics. 2050 climate change projections indicate a reduction in amaranth physiological response and the intervention of PV with varying transparencies showed potential in reversing these adverse effects. When agrivoltaic interventions were applied, 69% transparent c-Si modules mitigated most negative impacts of the 2050 climate scenario while enhancing vegetative growth and plant height compared to control plants (Fig. 4). Although 80% transparent Cd-Te PV modules also improved plant height to a similar degree, the detrimental effects of the 2050 climate scenario on physiological performance were still evident. The panel appeared to reduce the solar energy available to the plants more than those cultivated under the 69% transparent c-Si modules or no agrivoltaics, as indicated by lower PAR values and leaf temperatures. The severe reduction in light availability seemed to have decreased intracellular CO_2_ and limited Photosystem II efficiency. This reduction seems to be tied to low light instead of heat stress since these plants have better Photosystem II efficiency values and lower leaf temperatures than both 69% transparent c-Si and non agrivoltaic plants. This may be attributed to the uniform shading pattern offered by Cd-Te PV module compared to the non-uniform shading provided by its c-Si counterpart. These findings are similar to results from Jamil et al. (2025) ^25^ and highlights the importance of PV module transparency and material when optimizing an agrivoltaic system that can negate the effects of climate change in amaranth.

The 69% transparent c-Si module enhanced the physiological responses of amaranth when compared to plants cultivated with and without agrivoltaic intervention under 2050 climate conditions. The recorded PAR values for these plants were lower than control plants, but higher than those under 80% transparent Cd-Te PV. This balance optimized PAR which was reduced appropriately to counteract heat stress while allowing necessary light availability needed for Photosystem II activity. This was facilitated by open stomata, increasing intercellular CO_2_, stomatal conductance and transpiration rate. These benefits align with prior studies highlighting the ability of optimal PV shading in improving the physiological responses in different crops like jalapeños, tomatoes and chiltepin peppers ^39^. In developing sustainable and climate resilient agrivoltaic systems, emphasis must be placed on optimization using PV modules that are most beneficial for different kinds of crops.

### Assessing the effects of Agrivotaics production on amaranth growth performance and physiological response under current and future climates

Integrating both agronomic and physiological parameters using multivariate models like PLS-DA determined the variables responsible for plant performance in each climate scenario. In current climate conditions, leaf area, photosynthetic rate, fresh weight (week 5), stomatal conductance, intracellular CO□, and chlorophyll fluorescence were identified as vital in optimizing plant performance under agrivoltaics. Furthermore, PLS-DA also emphasized that plant height (week 5), minimum fluorescence, leaf temperature, and fresh weight (week 3) were variables that affected plant performance when cultivated under agrivoltaics in 2050 climate conditions. This identifies physiological variables and growth parameters that are imperative in optimizing amaranth performance under current agrivoltaic systems, while revealing the parameters susceptible to climate change when considering future agrivoltaic design that prioritizes amaranth health and production.

In addition, Pearson correlation analysis identified a strong negative correlation between plant height and minimum fluorescence suggesting that taller plants under 2050 conditions may have maintained more functional Photosystem II. Leaf temperatures were also positively correlated with both plant height and fluorescence stress, reinforcing the role of heat stress in photosynthetic inhibition ^40,41^. PLS-DA results also support findings in other crops, such as *Coffea arabica*, where internal CO□ and fluorescence traits indicated environmental stress responses ^42^. This demonstrates the broader utility of bioinformatics tools and its importance in assessing climate impacts and mitigation strategies when evaluating emerging crop production systems. The application of multiple multivariate approaches provided valuable insights into which PV types would be best suited to mitigate the adverse effects of climate change on amaranth.

### Economics Implications from Harnessing Agrivoltaic Food Systems

When considering incorporating agrivoltaic food systems, there are agricultural benefits as well as, economic potential. Agrivoltaic food systems improved plant yield and will also provide revenue from sustainable solar energy that in combination will surpass traditional farming.

## Conclusion

This research highlights the need for developing climate-resilient and sustainable food systems as climate change becomes threatening to global food security. Agrivoltaics presents a promising solution to maintaining and potentially improving future food production.

Additionally, this study provides an outline for enhancing food security in vulnerable regions across Canada, including remote, rural, coastal, and Indigenous communities. Furthermore, this work demonstrated that bioinformatics could be used to help understand the interactions between amaranth performance under changing climate and a decision support tool in the selection of the best PV modules that improve crop performance and be used to develop or innovate suitable climate resilient cropping systems. Future research should optimize the transparencies for amaranth production and the use of the 69% transparent c-Si modules and evaluate their effects on various leafy vegetable crops to expand agrivoltaic applications.

## Methods

### Experimental Setup and Data Collection

Historical climate data obtained from the Environmental Canada website ^43^ was used to simulate the current and future climates for crop growth. Amaranth (*Amaranthus Viridis*) was grown in biomes at the Biotron Institute for Experimental Climate Change Research Centre at the University of Western Ontario to assess their performance under current (2024) and projected (future-2025) climate conditions. Plants in Biome 2 were subjected to 2024 climate conditions, with an average daytime temperature of 25°C and an average nighttime temperature of 15°C and CO□ concentrations of 400 ppm. Biome 5 simulated projected 2050 climate conditions with an average daytime temperature of 29°C, an average nighttime temperature of 19°C, and CO□ concentrations of 550 ppm. Plants in both biomes were grown in a 16-hour daytime cycle and an 8-hour nighttime cycle. To see the effect of agrivoltaics as an innovative climate resilient system for amaranth production, the plants were also grown under PV modules of varying transparencies. Plant responses were assessed in the same two climate conditions under c-Si modules with 44% and 69% transparency, Cd-Te modules ranging from 10% to 80% transparency, and a 25% transparent c-Si panel which spectrally shifts light into red wavelengths ^44–46^.

For all treatments, plant height and the number of leaves were recorded weekly. The plants were harvested at 8, 10, 12, 14, and 15 weeks after planting. PAR readings were recorded at plant height under cloudy conditions. Leaf dimensions and plant height for each plant were also measured in cm. Additional variables, such as stomatal conductance, photosynthesis, cellular CO□, and transpiration rate, were recorded using the Li-Cor 6800 system. Chlorophyll fluorescence was assessed through measurements of the F0, FV, FM, and FV/F0 were collected using a Handheld Chlorophyll Fluorometer OS30P+ (V S Instrument Private Limited) for each plant. These variables were collected and stored in a Microsoft Excel 2010® sheet for later analysis.

### Data Pre-Processing

Before analysis, the datasets were organized and cleaned. Given their size and complexity, organization began with data ontology, where abbreviated variable names are standardized and documented to ensure consistency and ease of use. Data tables were restructured to include the averages of numerical variables such as plant height, fresh weight, and the number of leaves, recorded over the five weeks of plant growth. Missing values were addressed by imputing the mean of the corresponding experimental group. Datasets were screened visually using scatter plots and bar plots to examine data structure, distribution, variation, and outliers. Relationships between variables like average plant height, number of leaves, fresh weight, photosynthesis, and stomatal conductance and other variables across climate conditions were also visualized.

### Data Analysis

The data was transformed as part of a data normalization process then tested to ensure homogeneity, heteroscedasticity and normal distribution with low outliers ^47^. Normality of data distribution was assessed using Q-Q plots. After determining that it was normally distributed, homogenous and heteroscedastic, parametric tests like ANOVA and Pearson’s correlation analyses were applied where most appropriate. To evaluate how climate change may impact Amaranth production and quality, the following steps were undertaken: 1) An initial exploratory data analysis using scatter plots and bar plots were used to visualize the structure of the dataset. This step highlighted data spread, variation, outliers, missing values, and simple relationships between key variables such as average plant height, number of leaves, fresh weight, photosynthesis, and stomatal conductance across 2024 and 2050 climate conditions. 2) Several multivariate analyses were applied to identify associations between plant response and physiological variables under climate change. Techniques such AHC, k-means clustering, PCA, Factor Analysis, RDA, and PLS-DA were used to detect unique patterns and relationships within the dataset (Supplementary Figs. 1 & 2). These techniques were also used to reduce the dimensionality of the dataset, making it easier to interpret large and complex datasets by identifying patterns or clusters of the most important variables affecting crop performance. 3) Outputs from the assessment above indicate the data was normal. As such, univariate analyses including ANOVA and correlation analyses, were conducted on specific plant performance variables that clustered together during the multivariate analysis. This step assessed their statistical significance in contributing to plant performance under each climate condition. These statistical tests help determine which factors most influence plant growth and how they can be optimized under different climate conditions.

To determine how amaranth performance varies under PV modules of different transparencies in current and future climate conditions, the following steps were undertaken: 1) PV modules were first grouped by transparency and material type to facilitate analysis. Modules with transparency below 30%, between 30% and 60%, and above 60% were categorized as low, medium, and high transparency, respectively. This categorization was essential for understanding how different levels of shading or PV material impacted plant growth and physiological responses under both climate conditions. 2) The dataset underwent multivariate and clustering analyses to determine which approach best grouped variables and treatments of relevance based on the data structure. Given the complexity and high dimensionality of plant physiology data, several statistical techniques were applied, including AHC, k-means clustering, PCA, Factor Analysis, RDA, and PLS-DA. PCA helped to reduce data dimensionality by identifying patterns between plant growth, physiological variables, and PV transparency. PLS-DA classified variables to determine how the treatment group or climate in which Amaranth was grown influenced plant growth and physiology. RDA mapped variables in an ordination space to visualize how climate conditions affected plant growth and physiology. AHC and k-Means was used to categorize variables based on statistical similarities, forming natural groupings. These techniques helped identify the most influential factors affecting plant performance under agrivoltaic systems in each climate condition. 3) Since the data was parametric, univariate analyses like ANOVA and Pearson’s correlation analysis were used to assess the statistical significance of relationships identified in the multivariate and clustering analyses. These methods were used to highlight variables that significantly contributed to plant performance under different PV transparency levels and climate conditions. All analysis was conducted used XLSTATS Premium package (version 26.4.1, Addinsoft, Paris, France).

## Acknowledgements

The authors would like to thank Md. Motakabbir Rehman, S. Khan, and S. Rana for technical assistance and helpful discussions. This work was supported by the Natural Sciences and Engineering Research Council of Canada and the Thompson Endowment.

## Nomenclature

Photovoltaic cells (PV), Fresh Weight and week recorded (FW#), Plant Height and week recorded (PH#), Number of Leaves and week recorded (NL#), Leaf Temperature (Leaf Temp.), Photosynthesis (Photo.), Stomatal Conductance (Stomatal), Intercellular CO2 (CO2), Transpiration (Transp.), Minimum Fluorescence (F0), Maximum Fluorescence (FM), Variable Fluorescence (FV), Initial Fluorescence Response (FV/F0), Photosynthetic Efficiency (FV/FM), Photosynthetically Active Radiation (PAR).

