## Supplementary Data for "Integrating Amaranth and Agrivoltaics in the Creation and Assessment of Climate Resilient Food Systems"

**Nomenclature**: Photovoltaic cells (PV), Fresh Weight and week recorded (FW#), Plant Height and week recorded (PH#), Number of Leaves and week recorded (NL#), Leaf Temperature (Leaf Temp.), Photosynthesis (Photo.), Stomatal Conductance (Stomatal), Intercellular CO2 (CO2), Transpiration (Transp.), Minimum Fluorescence (F0), Maximum Fluorescence (FM), Variable Fluorescence (FV), Initial Fluorescence Response (FV/F0), Photosynthetic Efficiency (FV/FM), Photosynthetically Active Radiation (PAR).

**Supplementary Information**

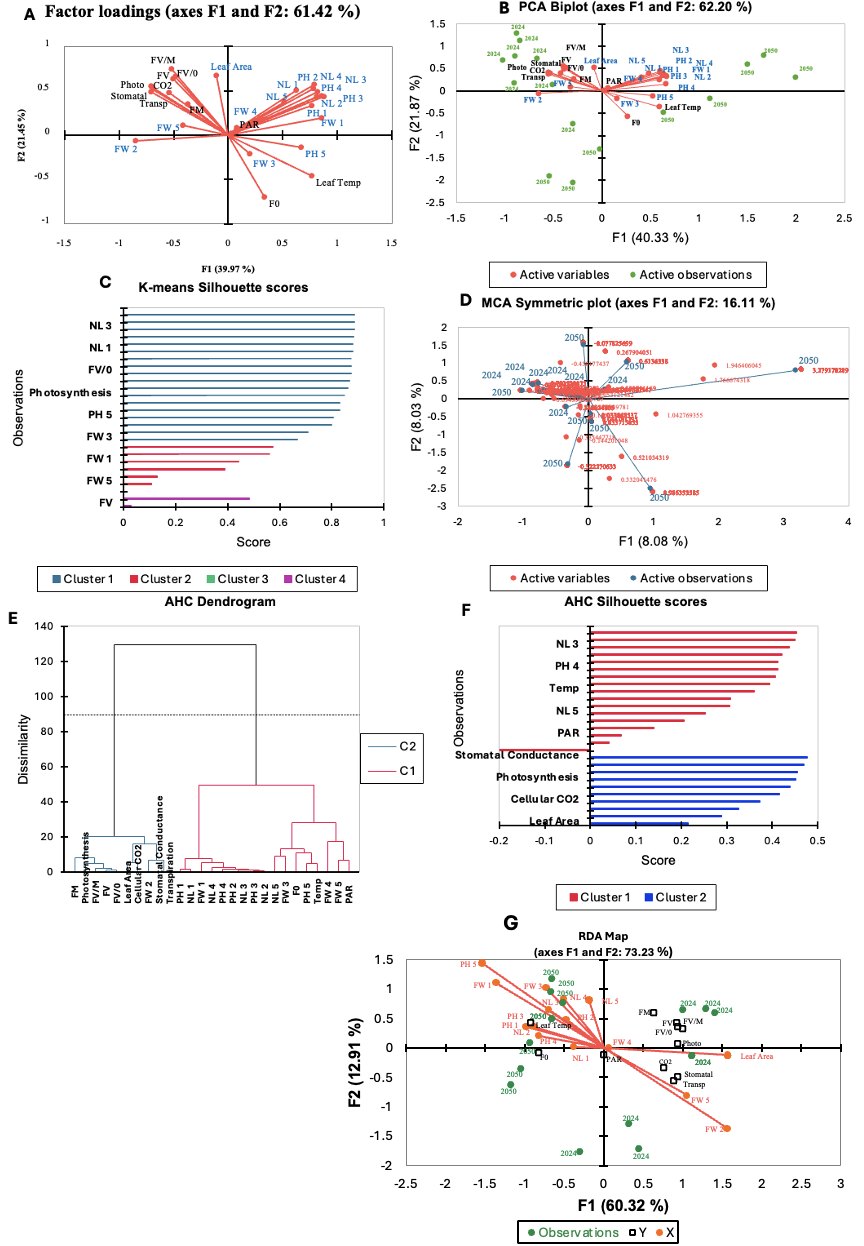

**Supplementary Figure 1. All multivariate analysis conducted to best assess the structure of the amaranth physiological and growth response data to simulated current and future climate when grown without PV.** Variables were clustered using Factor Analysis (A), PCA (B), k-means clustering (C), MCA (D) , AHC (E, F) and RDA (G). Factor Analysis, PCA and RDA had scores above the 60% threshold but did not capture the highest variability in the data (A, B, G). MCA, with a score below the 60% threshold and thus failed to account for significant data variation (D). k-means clustering identified three variables with a silhouette score below 0.6, indicating poor groupings (C). The AHC dendrogram categories variables into two distinct clusters (E), however, the silhouette scores for all variables remained below 0.6, suggesting weak data representation (F).

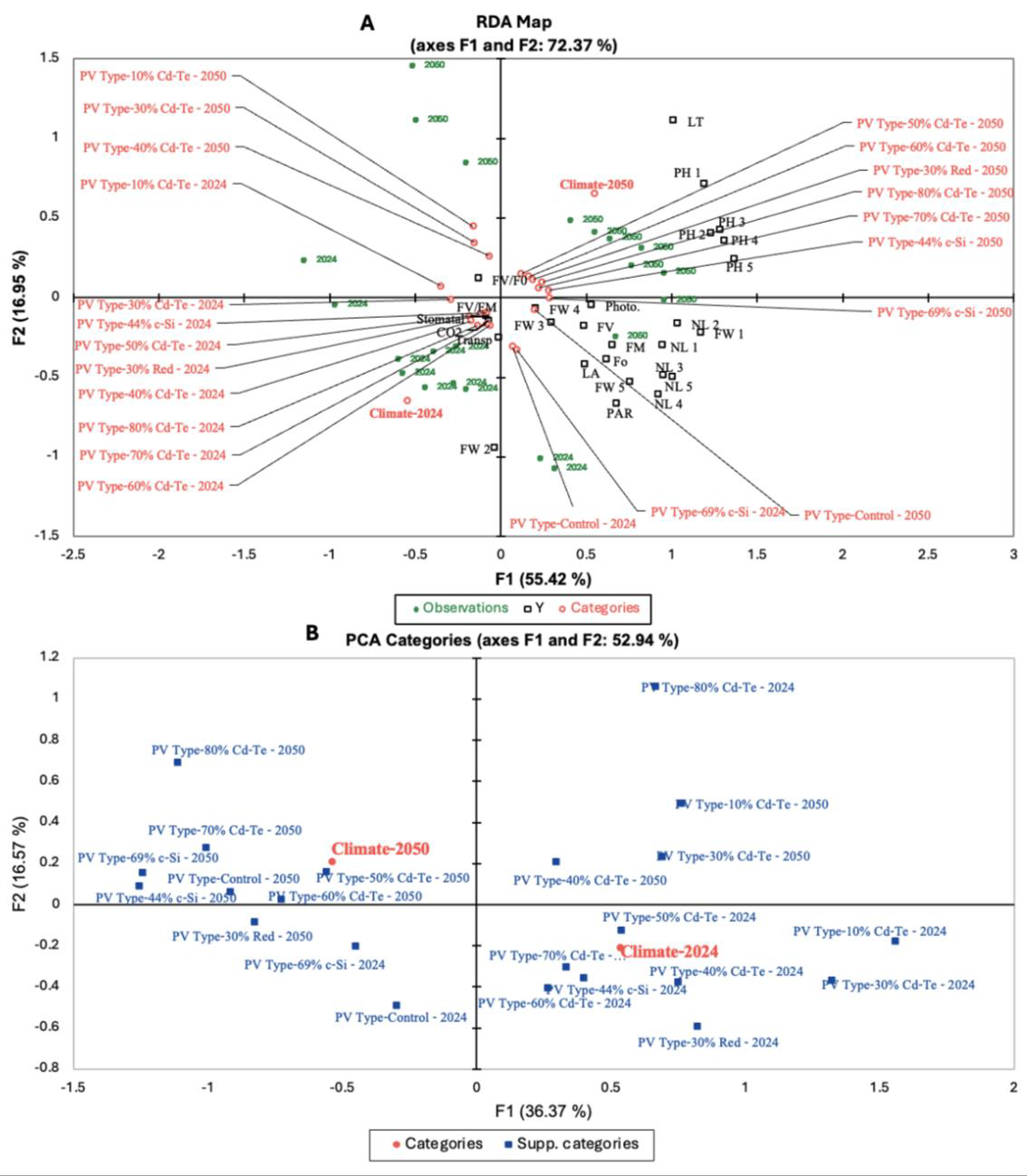
**Supplementary Figure 2. All multivariate analysis conducted to assess the best structure of the amaranth physiological and growth response data to simulated current and future climate when grown without PV.** Variables were clustered using RDA (A) and PCA (B). RDA achieved scores above the 60% threshold but did not capture the highest variability of the data (A). PCA , with a score below the 60% threshold , failed to capture significant data variation (B).

**Table S1. Physiological and growth parameters of plants grown under 69% transparent c-Si agrivoltaic treatment and without agrivoltaics in 2050 climate conditions.** Each parameter is represented as the mean ± SEM from n = 8 replicates per treatment. The percent change is denoted as difference to control and was calculated as ((Treatment − Control) / Control) × 100.

| **Parameter** | **Control (mean ± SEM )** | **69% c-Si PV (mean ± SEM)** | **Difference to Control (%)** |
| --- | --- | --- | --- |
| Plant Height | 34.25 ± 2.40 | 35.18 ± 1.72 | 2.72 |
| Number of Leaves | 39.88 ± 6.09 | 42.65 ± 1.99 | 8.24 |
| FV/FM | 0.64 ± 0.02 | 0.67 ± 0.02 | 4.69 |
| Stomatal Conductance | 0.06 ± 0.00 | 0.12 ± 0.00 | 100.00 |
| Intercellular CO_2_ | 153.23 ± 6.30 | 161.49 ± 7.79 | 5.39 |
| Transpiration Rate | 1.66 ± 0.01 | 2.87 ± 0.03 | 72.89 |
| Leaf Temperature | 30.43 ± 0.38 | 30.60 ± 0.08 | 0.59 |
| Photosynthetic Rate | 19.88 ± 0.48 | 22.29 ± 0.16 | 12.11 |
| Leaf Area | 117.70 ± 4.38 | 108.46 ± 12.0 | -7.85 |
| Total Yield* | 388.94 ± 8.72 | 380.92 ± 9.16 | -2.06 |

*The total yield during all weeks of harvest were calculated instead of the averages.
